# Shaping Sounds with P300 Based Brain-Computer Musical Interface

**DOI:** 10.1101/2022.06.13.495880

**Authors:** Uğur Can Akkaya, Çağatay Demirel, Reuben de Lautour, Gökhan İnce

## Abstract

This paper describes the development and testing of a brain-computer musical interface (BCMI) that allows a user to select and transform one element of a musical texture by paying attention to that particular element. In order to realize the BCMI system mentioned, a comprehensive testing scheme was established which uses auditory evoked potentials to elicit P300 waves via averaging various types of stimuli. Resented sound stimuli were divided into multi-channel speaker setups to have better localization of user-focused sound stimuli. A sound synthesis model was developed for transforming its sound texture based on neural oscillations that were categorized with the help of a self-organizing map algorithm. In addition, an artificial neural network was used to predict the possible P300 waves that show the attentional focus of a subject. Most of the P300 waves were classified successfully for most of the participants. Promising results were achieved concerning the developed BCMI system. A neural network model was also utilized to predict the possible P300 waves, which show the subject’s selective attention. The majority of the participants were able to correctly classify P300 waves. The proposed BCMI system yielded promising results.

## Introduction

The idea of using brainwaves for making music has been around since the 1960s with names like Alvin Lucier, David Rosenboom, and Richard Tieltelbaum (Brause, 2002). Technological advancements and the increased availability of bio-sensors such as electroencephalogram (EEG) have increased interest in the research field known as Brain Computer Musical Interfacing (BCMI) (Miranda, 2014). Such interest has resulted in a plethora of intriguing interactive systems that work primarily with brain waves (Christopher, Kapur, Arnegie, & Grimshaw, 2014). The BCMI field is an interdisciplinary research field which mainly feeds off computer science, neuroscience, and music research fields. By harnessing many aspects of these disciplines, it is possible to use different neuroscientific phenomena such as Event Related Potentials (ERP) (Luck, 2005). There are many prominent examples that use ERPs to make music, especially in Visually Evoked Potentials (VEP). In his article, Grierson (2008) mentions systems that use P300 components, such as P300 composer and P300 Scale Player. Another example would be Pinneger, Hiebel, Wriessnegger, and Müller-Putz’s (2017) BCMI system. In their work, they implemented a control system similar to the P300 speller, but instead of letters, music notations were used to control an open-source scoring system. Vamvakousis and Ramirez’s (2011) research is one of the P300 research that is closely related to this project’s motivation. In their study, Vamvakousis and Ramirez created a BCMI sequencer in which the melody that is being played can be influenced by focusing on or listening to the desired note in the sequenced melody.

The motivation of this work is to make a BCMI system where users of this interactive system can influence an element of the sonic texture around them only by listening to and focusing on that element. This system is an attempt to realize the possibilities of such an instrument. Building on previous works by Vamvakousis & Ramirez (2014, 2015), Käthner, Ruf, Pasqualotto, Braun, Birbaumer, & Halder (2013), Schreuder, Blankertz, & Tangermann (2010), and Biroschak, Kleschen, & Smith (2005), several new approaches and refinements were presented in this work. First, the authors investigate the potential of sound localization in a BCMI system to more effectively isolate individual sounds from a larger sonic texture through the use of a multichannel speaker array. Second, a multi-target oddball paradigm was designed to improve auditory evoked potentials (AEP) acquisition that will give better flexibility and musicality to the BCMI system by increasing the target stimulus choice for the subjects. Third, the system performs simultaneous real-time acquisition of two types of brain wave data. Target identification was enabled through the acquisition of the P300 wave of AEP while simultaneously recording neural oscillations from brain waves to enable meaningful transformation of synthesized sounds. P300 wave is an ERP that is considered to be an endogenous potential.

The determination of the P300 wave was realized using machine learning algorithms. Processing of neural oscillations for smooth transitions between changing parameters was supported by the self-organizing maps (SOM) algorithm.

## Methodology

The aexperiments were run in an acoustically sealed music studio. The tests were carried out on ten healthy individuals aged between 24 and 30 years old. Every participant gave their permission for their EEG data to be recorded and utilized in this study. The participants sat in the center of a five-channel array, equidistant from each speaker. Except for the two rear speakers, which played the same frequent stimulation, each speaker presented a separate sound stimulus. The display in front of the subject uses color-coded dots to show which speaker was playing a sound stimulus in real time. The sound stimuli had an interstimulus interval (ISI) of 400 milliseconds, with 300 milliseconds of sound stimuli and 100 milliseconds of silence. Before the experiment, each participant is given a brief presentation. While the EEG equipment records their neural responses, participants were instructed to concentrate on their preferred target stimulus and mentally count the occurrences of that sound stimuli in order to affect the sound. The P300 component was determined using ANN using real-time EEG data. The experiment was split into two parts in order to train the ANN model. The respondents were instructed to focus on predetermined target stimuli during the first three to six runs. Afterwards, the ANN model was trained using the focused target stimulus as well as the other two non-target stimuli from each session. The training trials were kept to a minimum in order to avoid exhausting the individuals. Participants in the training trials received no sound changes (audio-visual feedback). The second portion of the experiment comprised a real-time scenario in which subjects were asked to choose one auditory stimulus among three. The parameters of the synthesizer responsible for producing that sound were altered based on the brain oscillations classified by the SOM if the ANN anticipated a likely P300 wave. Fig. 1. depicts a flowchart of the system overview.

**Fig. 1.**
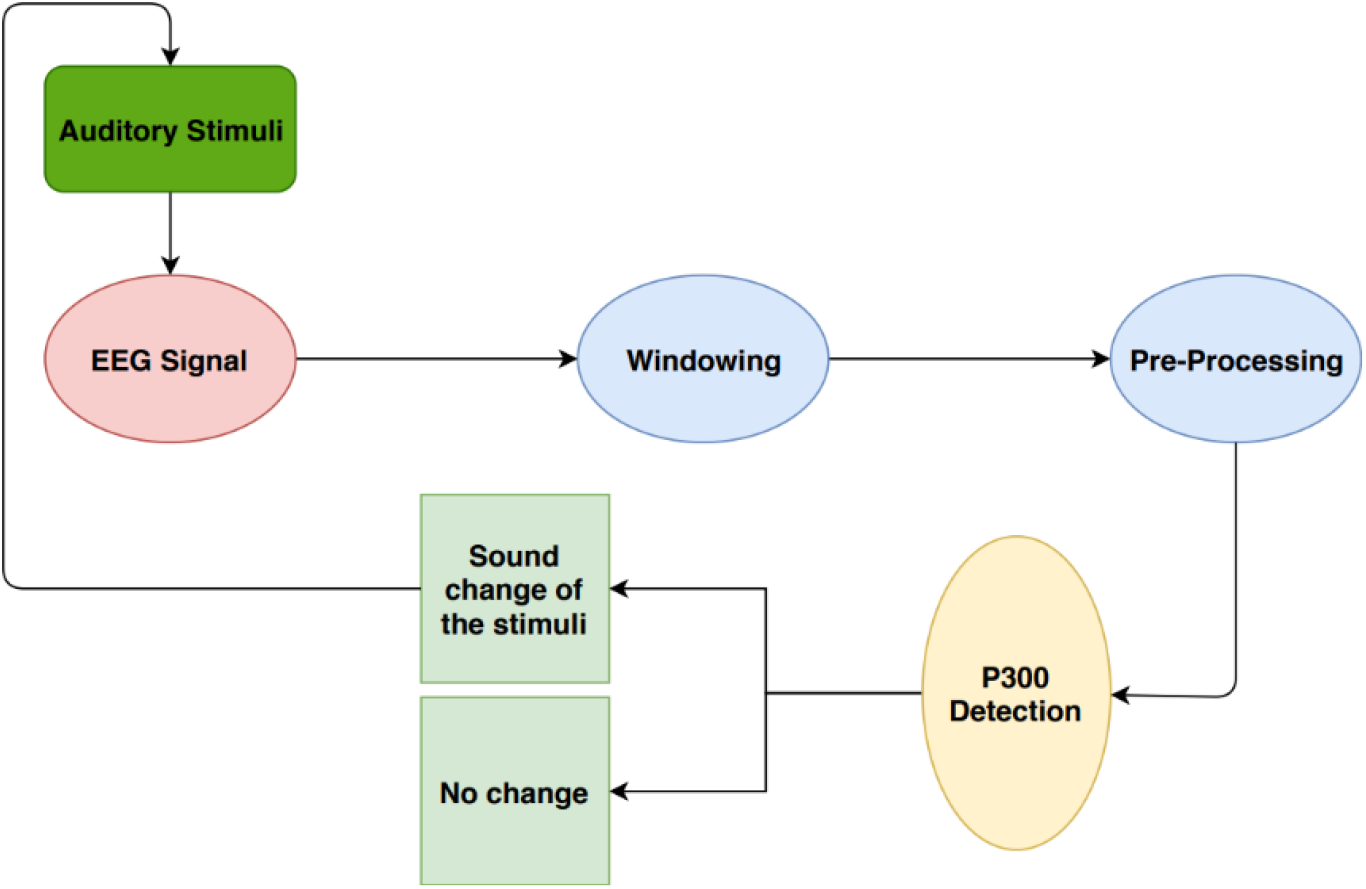
System overview.

## Sound Localization and Probability Distribution for Speaker Configuration

Sound localization was applied to increase the subject’s focus on the stimulus they had chosen. A saw-tooth wave between 1000 and 2500 Hz was employed to help the subjects better localize the sound stimulus. Aside from the two back speakers, which played a 1000 Hz saw-tooth wave, each speaker produced a separate individual frequency. The experiment setup is shown in Fig. 2.

**Fig. 2.**
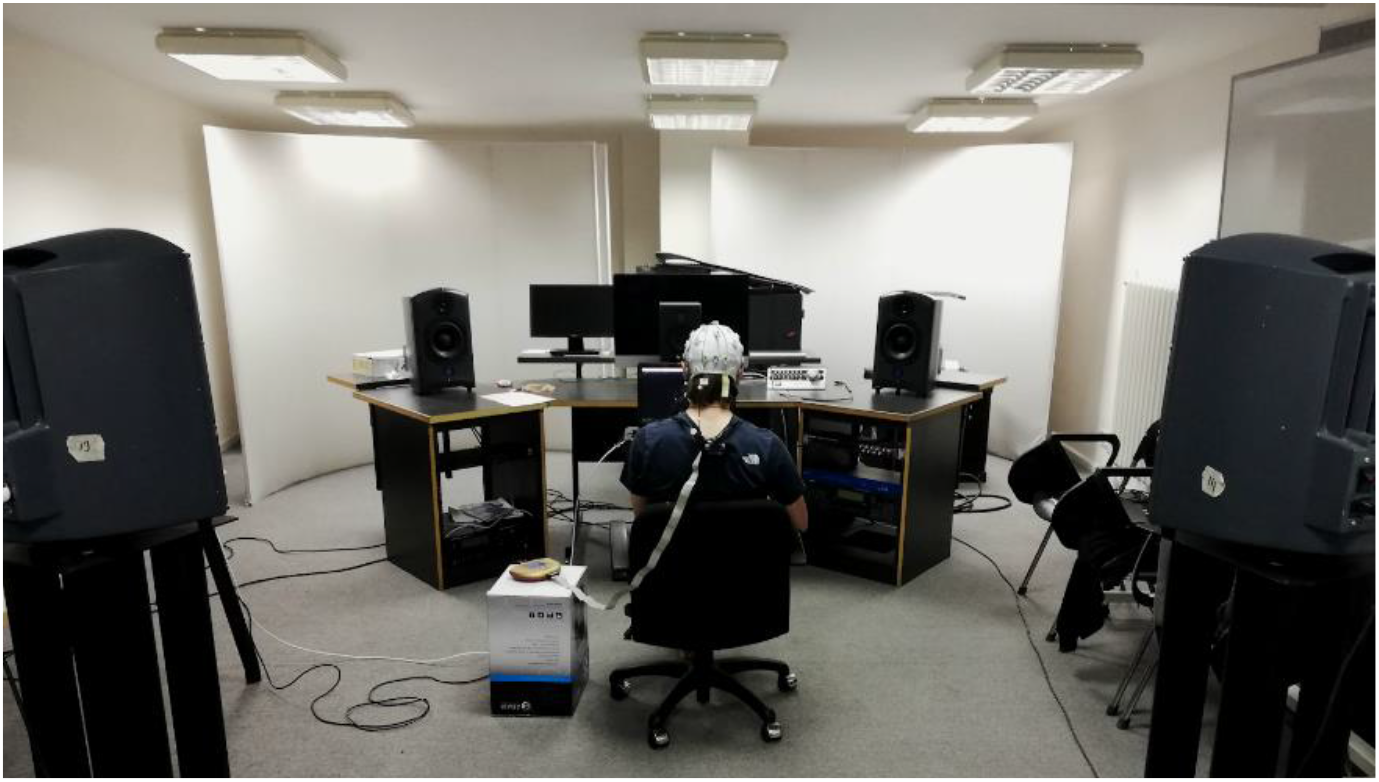
Experimental setup.

## Using Neural Oscillations as Synthesis Parameters

Neural oscillations are neuroscientific phenomena that can be observed in certain frequency windows of the raw EEG recording. These rhythmical neural patterns are related to the cognitive and bodily functions of the participant. These patterns contribute to many cognitive functions (Başar, Başar-Eroğlu, Karakaş, & Schurmann, 2001). There are five main types of neural oscillations: delta, theta, alpha, beta, and gamma (Liu, Chiang, & Chu, 2013). Table 1 shows the neural oscillation types.

**Table. 1.** Neural oscillation types.

| Activity | State |
| --- | --- |
| Delta (0.5 – 3 Hz) | Deep sleep, slow-wave activity |
| Theta (3 – 7 Hz) | Relaxation, low alertness, or daydreaming |
| Alpha (8 – 12 Hz) | Resting, calmness and mindfulness |
| Beta (12 – 30 Hz) | Attentive, thinking and focusing |
| Gamma (30+ Hz) | High alertness and conscious feeling |

A sound synthesis technique that incorporates intelligible data acquired from brain waves was developed inside the custom-created Max Msp patch. A subtractive sound synthesis model was used for the sound stimuli. The fundamental value for each auditory stimulus was determined before the experiments. The created Max Msp patch multiplies the recorded neural oscillation value (μV square per Hz) by the fundamental values of auditory stimuli to create upper harmonics to the synthesized sound. Real time segmented neural oscillations were used to alter the balance of the partials of the determined fundamental frequency of the synthesizer. Each parameter that affects the upper harmonics of the auditory stimulus that was determined as P300 true by the trained ANN model, changes in relation to the 600 milliseconds of EEG data recorded before the onset of the instantaneous target stimulus. It should be noted that sound change and sound morphing only applied to the target stimulus that was marked as a P300 true candidate. Other auditory stimuli were not affected by this process unless the user decides to change their desired focused stimulus.

Neural oscillations recorded from the segmented raw EEG data were observed to be disjunct. The reason for these big jumps was that each segmented EEG data consisted of relatively different neural oscillations related to the stimulus that came before it. In order to eliminate this disjunctive data, the SOM categorization algorithm was used as an unsupervised categorization method. The recorded neural oscillations were fed to the SOM for categorization of neural oscillations. The output of this algorithm was used for smooth parameter changes of the subtractive synthesizer created.

## Preprocessing and P300 Classification

Data acquisition was performed with a Brain Products V-amp 16 EEG device, using Fp1, Fz, Cz, and Pz channels. A custom python script was created for signal processing techniques used on the raw EEG data.

Preprocessing techniques were used to eliminate noise factors that occur due to the low SNR ratio of the EEG recordings (Luck 2015). After each sound stimulus, 600 milliseconds of EEG signal were segmented and recorded. Preprocessing was performed on each segment of recorded EEG data. Windows were band-pass filtered between 0.1 and 30 Hz. A 50Hz notch filter was used to eliminate DC hum. Baseline correction was implemented to reduce drifts in the raw EEG. A threshold of 70 μV used for artifact rejection. This process is necessary for eliminating blinks, ocular movements, swallowing, and body movements that can alter the P300 component analysis (H. Mirghasemi, M. B. Shamsollahi & R. Fazel-Rezai). The identified trial window was excluded from the averaging process if the artifact rejection algorithm finds any noise inside a segmented window, which can occur due to bodily and ocular movements.

P300 classification was tested using four different machine learning techniques. Support vector machines (SVM), linear discriminant analysis with shrinkage value (LDA), random forest, and ANN were examples of these techniques. Previous researches have shown that LDA with shrinkage value enhances ERP classifications, and that LDA outperforms other classifiers in a dataset containing target and non-target P300 signals from diverse electrode positions (Höhne, Blankertz, Müller, & Bartz, 2014; Mirghasemi, Fazel-Rezai and Shamsollahi, 2006). Furthermore, SVM is a well-known accurate classifier for P300 research (Krusienski et al., 2006; Thulasidas, Guan, & Wu, 2006). ANN variations have been used in various sub-fields of neuroscience to solve complex problems such as pattern recognition, speech, vision, and dynamic data processing (Jaiyen, Lursinsap & Phimoltares, 2010; Washizawa, 2010; Ozawa, Roy & Roussinov, 2009; Ge S.S., Yang Y. & Lee T. H., 2008).

The offline succession metric for each EEG channel in each subject’s dataset was 10-fold cross validation, and it is proven that the channel locations chosen for classification results varied. In all the trials, Fz channel was shown to be the most relevant channel for estimating target P300 signals among chosen scalp electrodes, because the dataset derived from Fz yields the best classification results in all classifiers on the collected dataset. Each machine learning algorithm undergoes hyperparameter optimization, and multiple combinations of hyperparameters within each model were trained independently, a process called grid search (Bergstra, Komer, Eliasmith, Yamins & Cox, 2015). The ANN model outperformed the other three machine learning algorithms in cross-validation accuracy out of four different classification methods. The chosen ANN model contains two layers, comprising 80 neurons in the first layer and 80 neurons with a ReLu activation function in the second layer. With a learning rate of 0.001 and a momentum value of 0.9, Adam was picked as the optimizer. A total of 50 epochs were used to train the final model.

The recorded data from the training stage of the experiment was utilized to train the ANN model in order to build the dataset required to classify the P300 response. Each subject’s dataset undergoes subject-specific training. There were five different types of consecutive windows for the P300 wave classification, which were classified based on the number of windows utilized in the averaging procedure. There were 5, 10, 15, 20, and 25 windows in total. The cross-validation performance of these five windowing types was investigated. It was discovered that an average of 15 consecutive windows produced greater cross-validation accuracy from the P300 candidate data sets than an average of 5 or 10 consecutive windows for all window counts. For the P300 classification accuracy, an average of 25 consecutive windows does not yield the desired significance. Furthermore, classifying the P300 wave with a 25 window count takes longer since each auditory stimulus requires 25 consecutive windows to be filled. As a result, the mean values of each of the 15 preprocessed EEG windows were taken as a final dataset and labelled according to stimulus information. Based on the EEG data of participants’ focused stimuli, these averaged windows were input into ANN as target or non-target trials. The construction of a P300 candidate based on the oddball paradigm is the name for this procedure (John & Catherina, 1997). As a result, the dataset needed for concurrent P300 classification was ready for testing. P300 candidates of each stimulus type were assessed for each time step in which a stimulus occurs. For each P300 candidate, the probabilistic outputs of target stimulus estimation findings were examined, and the stimulus belonging to the highest estimated P300 candidate was accepted as a target stimulus.

## Experiment Result

The mean P300 classification accuracy for all subjects was found to be 88.01% for offline situations. which shows a positive result for the usability of the proposed BCMI system. The mean of real-time classification results was found to be 64,75%. The subjects generally reported that they felt able to influence the sounds. Fig. 3 shows one of the successive sessions where the user was focused on the stimulus labelled as type 1.

**Fig. 3.**
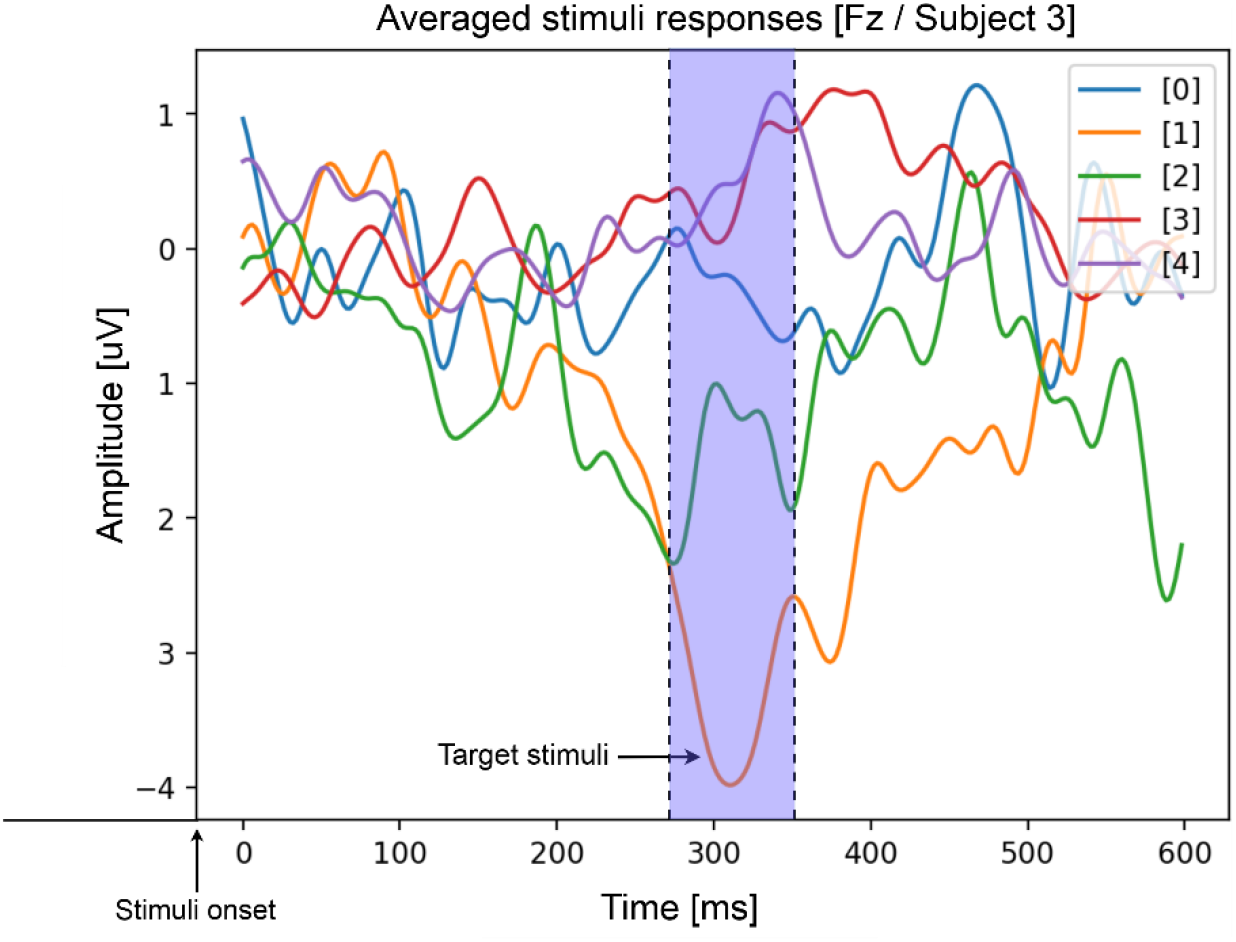
Example comparison between target and non-target stimuli responses (averaged windows per stimuli type) of subject 3. Expected amplitude desynchronization occurs around 300 ms after stimulus onset in target stimuli (labelled as orange/stimuli number 1).

However, not all trials were successful. This could point out the challenges of this kind of BCMI system where subjects’ focus and motivation are one of the important aspects that affect the P300 wave. Another important aspect would be the restricted body and ocular movements of the participant. Any change in posture can induce noise in the EEG recordings. In addition, three target stimuli could impose a problem because of the lower probability of deviant stimuli (0.0667%). A low probability could mean a longer wait for the same stimulus to play again. While this enhances focus on the chosen stimulus, at the same time, low repeatability could break a participant’s focus on the target stimulus because of the same unique appearance of non-target stimuli. Having said that, because of the high accuracy of the prediction of the P300 wave, subjective probability calculation for the multiple target oddball paradigm showed promising results. Evidently, probability calculation had a significant effect on the P300 wave.

## Conclusion

Experimental results showed that it is possible to use AEP in a multiple-target oddball paradigm with the help of subjective probability and sound localization. The model prediction performance was also promising, and the authors speculate that using ANN in similar BCMI systems might give usable results since the classifier was searching for possible P300 waves at the same time from each possible target stimulus. Through the use of subjective probability and sound localization, experimental results indicated that AEP may be used in a multiple-target oddball paradigm. The model prediction performance was similarly encouraging, and the authors suggest that utilizing a P300 predictor in comparable BCMI systems should yield practical results because the classifier searches for plausible P300 waves from each conceivable target stimulus simultaneously.

This study allowed authors to access information about the conditions under which such a system should be carried out. There are some limiting aspects to these conditions, such as low followability of the randomly distributed auditory stimuli and restricted body and ocular movements. Because of these constraints, developing a definitive BCMI system is considered to be difficult. Furthermore, because it employs advanced paradigms such as the multiple target stimuli oddball paradigm with surround ques for localization, this system necessitates a high level of attentiveness from its users. These requirements must be completed in order for the BCMI system to be successful. As a result, powerful computational signal processing techniques are essential to limit these restrictive and mandatory criteria, resulting in improved user accessibility.

One of the study concerns arising from the trials is the effect of sound stimulus type on the P300 component. Another fascinating topic to investigate further is deep learning algorithms for reducing the expected time for categorization of the successful P300 wave. This could tell a BCMI system which sound stimulus a subject is focusing on in real time. Another major component of the proposed BCMI system that can be improved is its compatibility with consumer-grade EEG equipment, which is much easier to set up and operate.

Building a BCMI system that allows users to affect and change sound texture just by focusing on a sound object inside that texture is an intriguing topic that deserves further attention. Because a system such as this could store a lot more information about human auditory perception, it might lead to greater use of our brainwaves in music production and a lot more aesthetic performances.

